# Effect of Artificial Lung Fiber Bundle Geometric Design on Micro- and Macro-scale Clot Formation

**DOI:** 10.1101/2024.01.05.574443

**Authors:** Angela Lai, Natsuha Omori, Julia E. Napolitano, James F. Antaki, Keith E. Cook

## Abstract

The hollow fiber membrane bundle is the functional component of artificial lungs, transferring oxygen and carbon dioxide to and from the blood. It is also the primary location of blood clot formation and propagation in these devices. The geometric design of fiber bundles is defined by a narrow range of parameters that determine gas exchange efficiency and blood flow resistance, such as fiber packing density, path length, and frontal area. However, these parameters also affect thrombosis. This study investigated the effect of these parameters on clot formation using 3-D printed flow chambers that mimic the geometry and blood flow patterns of fiber bundles. Hollow fibers were represented by an array of vertical micro-rods (380 micron diameter) arranged with varying packing densities (40, 50, and 60%) and path lengths (2 and 4 cm). Blood was pumped through the device corresponding to three mean blood flow velocities (16, 20, and 25 cm/min). Results showed that (1) clot formation decreases dramatically with decreasing packing density and increasing blood flow velocity, (2) clot formation at the outlet of fiber bundle enhances deposition upstream, and consequently (3) greater path length provides more clot-free fiber surface area for gas exchange than a shorter path length. These results can be used to create less thrombogenic, more efficient artificial lung designs.

**Translational Impact Sentence:** Fiber bundle parameters, such as decreased packing density, increased blood flow velocity, and a longer path length, can be used to design a less thrombogenic, more efficient artificial lung to extend functionality.

## 1. Introduction

Thrombus formation at the blood-material interface of medical devices has been an ongoing concern for more than 60 years.^6^ Of medical devices on the market, artificial lungs have one of the greatest blood contacting surface areas: ranging from 0.8 - 1.8 m^2^. The fiber bundles are densely packed for maximum gas exchanging efficiency, with blood flow external to the fibers and a mix of oxygen flowing within the fiber lumen. The blood-contacting surfaces of the fiber bundles experience almost immediate protein adsorption, leading to direct activation of coagulation cascade factors and platelet adhesion and activation. Coagulation cascade and platelet activation, in turn, lead to the formation of solid clot within the fiber bundle, thereby reducing the gas exchange efficiency and increasing blood flow resistance. Furthermore, embolization of clots may result in thromboembolic occlusion of blood vessels within major organs. To prevent this, patients are prescribed a variety of systemic anticoagulant drugs, which in turn cause increased risk of bleeding and thus must be used with caution. Despite best efforts to mitigate thrombosis, typical clinical artificial lungs clot and fail within 1-3 weeks.^1–8^

The fluid dynamic engineering of artificial lungs has long been used as a means to reduce clot formation by avoiding areas of blood stagnation or recirculation that are known to lead to accelerated clot formation. This design process has focused primarily on the inflow and outflow manifolds that direct blood to and from the fiber bundle. There is currently no proven relationship between the geometric design of the fiber bundle shape and fiber packing density and the speed of clot formation, despite this being the site of the most rapid clot formation. It is hypothesized that these fiber bundle parameters could be optimized to reduce thrombus formation and increase the useful lifetime of these devices.

There is also no current standard artificial lung fiber bundle geometry. Some fiber bundles are rectangular, such as the Maquet (Getinge) Quadrox, while others are cylindrical, such as the LivaNova EOS. Despite this, the shape and dimension of these fiber bundles can be simplified to the following parameters: (1) the path length (L), the distance from the inlet face of the fiber bundle to the outlet in the direction of blood flow, (2) the frontal area (A_f_), the area of the fiber bundle perpendicular to direction of blood flow; and (3) the packing density, the percentage of the fiber bundle volume that is occupied by the hollow fibers. Commercially available oxygenator fiber bundles utilize a wide range of these parameters, and there is no consensus on what makes the least thrombogenic fiber bundle from clincal data or computational simulations.^9–11^

In this experimental study, microfluidic flow chambers that mimic the local fluid mechanics within the fiber bundles of commercial oxygenators were constructed. The geometry of the chambers was varied to simulate a range of fiber bundle packing density, path length, and frontal area. Freshly drawn, lightly heparinized, human whole blood was pumped through these microfluidic chambers, and multi-scale clot formation was examined using confocal microscopy to map local platelet and fibrin deposition. Micro computed tomography (µCT) was employed to reconstruct the gross, 3D clot geometry.

## 2. Materials and Methods

### 2.1 Experimental Design of the Flow Chamber

To simulate the local conditions of an artificial lung hollow fiber bundle, miniature flow chambers were geometrically modeled using parametric computer-aided design software (Solidworks, MA) to represent a small section of the fiber bundle. These flow chambers were 7.3 mm wide by 3 mm tall and had varied combinations of the physical fiber bundle parameters, packing density and path length. All fibers were simulated as 380 µm cylinders (rods) oriented perpendicular to the flow direction. These rods were arrayed evenly spaced to create packing densities of 40, 50, or 60% (Figure 1). These rods spanned the entire 3 mm height of the chamber to create blood flow patterns similar to the hollow fibers of an artificial lung. The length of the flow chamber was also varied to examine the effect of path length on clot formation. Lastly, the blood flow velocity was varied to simulate the effect of the fiber bundle frontal area on the local fluid mechanics: inasmuch as larger frontal area reduces average blood flow velocities. Thus, the blood flow velocities were set at 16, 20, and 25 cm/min to simulate, for example, 2 L/min of blood flow through a full sized adult artificial lung with 80, 100, and 120 cm^2^ frontal areas, or 4 L/min through frontal areas half that size.

**Figure 1.**
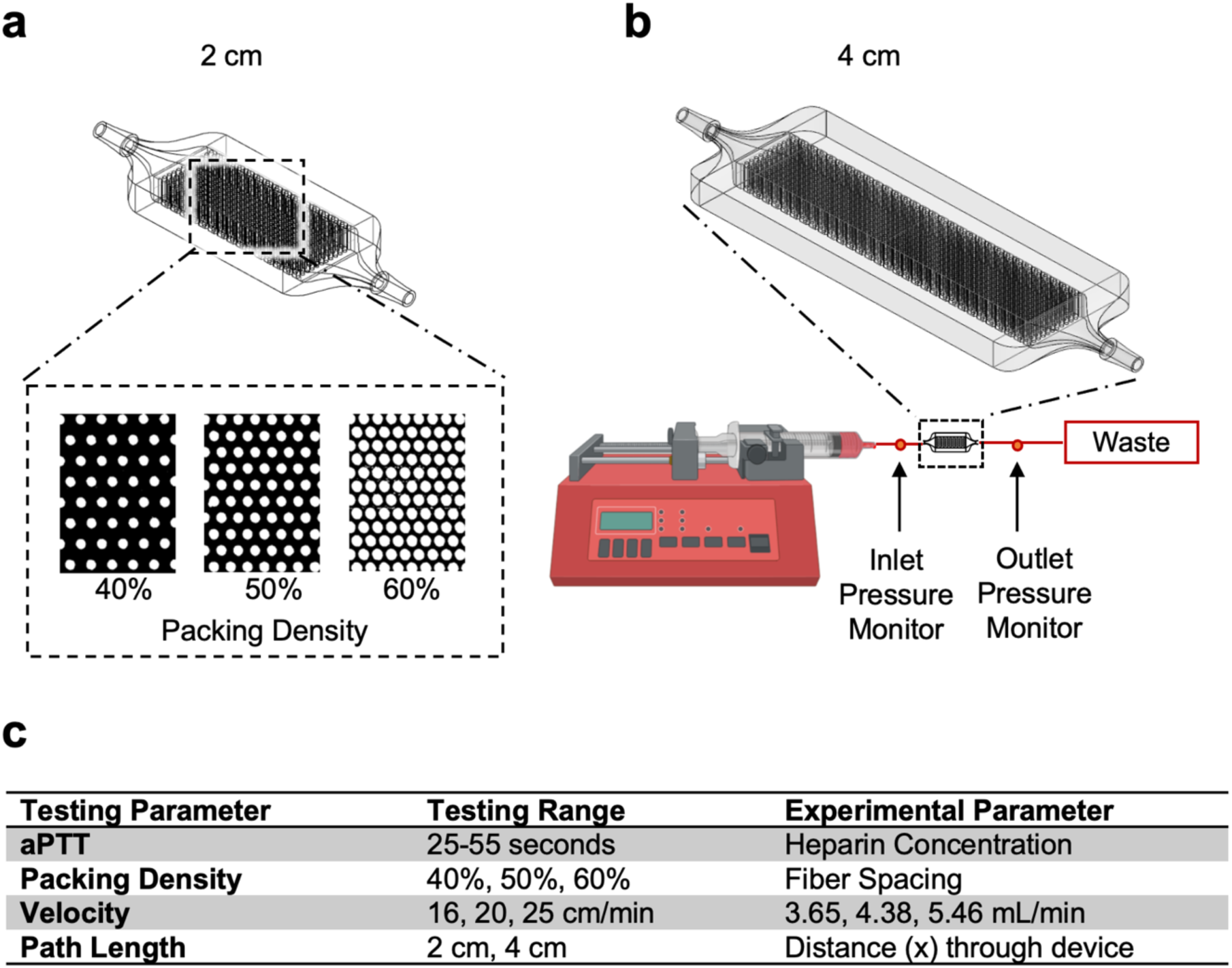
(a) The experimental flow chamber was designed to be 2 or 4 cm and with a packing density of 60%, 50% or 40% rods to simulate the flow path of blood through an artificial lung hollow fiber bundle. (b) The flow circuit involved one-way flow from a syringe pump, connected to the flow chamber and then to a waste container. A manometer measured the pressure drop across each experimental device. (c) The tested parameters in the benchtop experiments covered the range seen in clinically used oxygenators.

Each chamber was 3D-printed (Ember Autodesk, San Rafael, CA) using clear acrylate resin (PR-48, Colorado Polymer Solutions, CO) similar to urethane acrylate plastic for ease of repeatability and uniformity. Uncured resin was rinsed out using 60 mL of isopropyl alcohol, post-cured with a UV lamp for 20 minutes, and then sterilized using UV-ozone for 15 minutes.

Each of the independent design parameters (packing density, velocity, path length) was investigated by perturbing a baseline design with 50% packing density, 20 cm/min blood velocity, and 2 cm path length (Figure 1). As described previously, the packing density was adjusted to either 40 or 60%; the velocity was tested at 16 and 26 cm/min; and the path length was tested at 2 and 4 cm.

### 2.2 Experimental Evaluation of Microchannel Devices

The estimated, clot-free, baseline resistance and Reynold’s number was measured for each type of device using a glycerol-water mixture. During whole blood experiments, clot formed instantaneously upon blood contacting the device surfaces. Thus, a clot-free reference resistance was needed to effectively interpret the role of clot in increasing device resistance during whole blood studies. The glycerol-water mixture was titrated to a 3.4 cP viscosity using a viscometer (9721-B56, Cannon-Fenske, USA) and was pumped through at a rate of 3.65, 4.38, and 5.46 mL/min using a syringe pump (NE-300, New Era Pump Systems, Inc., NY) to achieve an average velocity of 16, 20, and 25 cm/min through the simulated fiber bundle. Pressure drop was measured with a manometer (CR410, Ehdis, China), and the resistance was calculated using the standard formula:

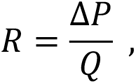

in which *R* is resistance, *ΔP* is the pressure drop across the device, and *Q* was the flow rate. Pressure measurements were made in triplicate for each flow rate and averaged.

Blood studies were conducted using human blood from consenting donors under an approved Institutional Review Board protocol. Between 250-550 mL blood was drawn from each participant with an 18 gauge catheter, as per standard venipuncture procedures using a gravity drain. The blood was anticoagulated with 0.1 U/mL of heparin. After the full volume of blood had been collected in a sterile blood bag, it was transferred to 60 mL syringes for the experiment within 10 minutes after the blood draw. A syringe pump (NE-300, New Era Pump Systems, Inc., NY) was used to infuse blood through the device in a single pass through 1/16” Tygon tubing (E-3603, Saint-Gobain, Malvern, PA), as seen in Figure 1. To minimize the effect of donor-to-donor variability, each experimental run was performed with all six configurations of flow chambers (three packing densities and two path lengths) utilizing blood from the same donor.

Differential pressure across the device was measured every 3 minutes over a total of 15 minutes using a manometer (CR410, Ehdis, China), and then used to calculate resistance using the equation above. Blood samples were collected at the inlet and outlet for measuring blood cell count (2 mL in K_2_EDTA) and aPTT/PT (3 mL in 1:9 citrate) using a clinical hematology analyzer (Diagnostica Stago Start 4, Siemens, Germany).^12^ If aPTT was outside the range of 20-50 seconds, the experimental data was not used. After 15 minutes of flow, the pump was stopped, the device was detached from the circuit and immediately gently rinsed with heparinized saline (2 U/mL) to halt the progression of clot formation. The flow chamber was then filled with 4% (v/v) paraformaldehyde and fixed overnight at 4°C.

### 2.3 Micro-Computed Tomography Scans

Micro-Computed Tomography (µCT) scans were performed to quantify the clot volume inside the flow chambers. Prior to the µCT scan, the fixative inside the flow chambers was flushed using 10 mL of saline. Three flow chambers at a time were placed inside a custom-built holder and then scanned using a Skyscan 1172 (Bruker-Skyscan, Contich, Belgium) µCT system that has a 3 µm voxel resolution and the following conditions: 40 Vp, 175 µA, 300 ms exposure, 0.4 degrees rotation step, 8 frame averaging. Serial scans were reconstructed into a 3D volume using NRECON (Bruker-Skyscan, Contich, Belgium) with a 60% beam hardening correction and a ring artifact correction of 20. These volumes were then converted to ND2 files (FIJI, ImageJ 1.51H, NIH Bethesda, MD) and imported into an open source software package (Slicer 3D 4.10.2) to be transformed, segmented, and cropped. The volume was cropped 0.4 mm from the top and bottom, 1.3 mm from each side, and 1 mm from the inlet side to avoid the edge effects induced by blood activation from the wall. The cropped volume was then imported as a DICOM image back into FIJI.

Thereafter, all of the images (n = 6) for each experimental condition were averaged to create a 3D map demonstrating the probability of clot formation at any voxel in the device. To isolate the clot volume from the background, three-dimensional reconstructions were created from scans of four clean chambers for each packing density and path length for a total of four masks, one for each testing parameter. These masks were then subtracted from each individual experimental scan, so the volumes analyzed only consisted of clot (Figure 2).

**Figure 2.**
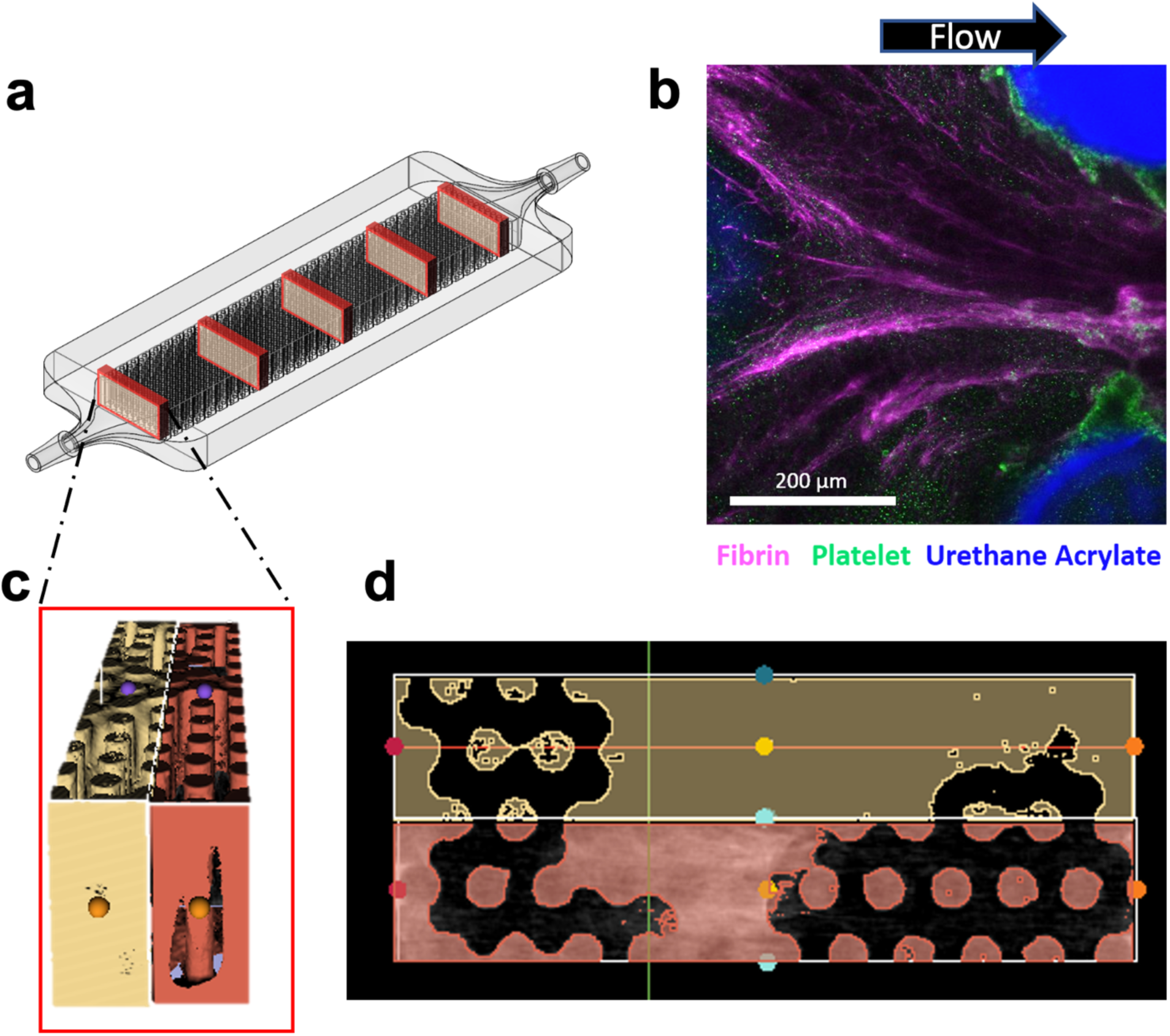
MicroCT or fluorescent microscopy was used to image the entire chamber. (a) For analysis of microCT images, five transverse slices were taken from the inlet to outlet and the volume was quantified. (b) For microscale analysis, fibrin, platelets were fluorescently labeled with antibodies to visualize deposition around the rods. (c) MicroCT slices were reconstructed to quantify the solid volume, consisting of clot and rods. (d) These 3-dimensional volumes were averaged across the Z-axis to form an averaged image to present as 2-dimensional representation of the clot. Image masks were then used to remove rods so that only clot remains.

Micro CT volumetric raw clot data was presented in three ways. Firstly, the resulting 3D volume was sliced transversely from the inlet to the outlet and the clot at each slice was extracted using a custom macro (Figure 2). Clot volume at 5 locations through the length of the chamber were summed for 2 mm segments and then graphed to quantify the deposition of thrombus in relation to the flow path in the chambers. Results were normalized based on the available volume for clot to form in each type of chamber using the formula:

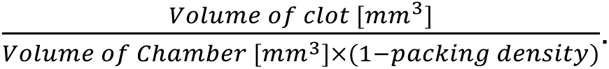

As a second means of normalization to account for differences due to surface-initiated blood activation, clot formation was measured at a specific distance from the inlet in each flow chamber. This distance was a function of the blood flow rate and packing density to find an equivalent blood-contacting surface area. This equivalent distance allowed for the clot volume to be compared for a set of conditions that would provide similar gas exchange due to the same surface area. For example, for devices run at 16, 20, and 24 cm/min, clot volume was compared at 6.8, 5.7, and 4.5 mm from the inlet, respectively, where each device had encountered the same fiber surface area. For the 40%, 50%, and 60% packing density devices, clot was compared at 10.0, 5.7, and 3.2 mm from the inlet, respectively.

Thirdly, for ease of display in a 2D image presentation, the 3D volume reconstructions of clot were averaged in the z-axis. This resulted in a top-down view of the probability distribution of blood clot deposited in the flow chamber for a clearer understanding of the spatial distribution of thrombus.

### 2.4 Immunofluorescent Staining and Imaging of Fibrin and Platelet Deposition

Additional experiments were performed to obtain microscale understanding of the clot composition within the microchannel device via platelet and fibrin fluorescent staining. The experiments were exactly the same as those in the previous section with the substitution of a glass coverslip on one side of the flow chamber for visualization.

In these experiments, packing densities of 40% and 60% and velocities of 16 cm/min and 25 cm/min were compared. After the 15-minute experiment, the flow chambers were rinsed with 10 mL of 2 U/mL heparin-saline and fixed overnight in 4% (v/v) paraformaldehyde. This was followed by six rinses with 1 X PBS. Blocking was conducted with 1% (w/v) bovine serum albumin and 5% (v/v) goat serum in PBS for 2 hours at 4°C.

For platelet and fibrin staining, 1:400 diluted CD61 stain (Tyr773 44-876G, ThermoFisher, USA) and a mouse monoclonal anti-human fibrin and fibrinogen (F9902, Sigma-Aldrich, USA) were incubated inside the flow chambers overnight at 4°C. The following day, after six rinses with 1 X PBS, the flow chambers were subsequently incubated with the secondary antibodies, goat anti-rabbit IgG (H+L) Alexa Fluor 555 (A-21428, ThermoFisher, USA) and goat anti-mouse IgG1 CF633 (SAB4600335, Sigma-Aldrich, USA), in a 1:250 ratio for 1 hour at room temperature. These flow chambers were then rinsed with 1 X PBS three times and imaged on a confocal microscope (A1 R+ HD25 Nikon) using a 16 X NA water immersion objective.

### 2.5 Statistical Analysis

All µCT clot volume and resistance values were analyzed for statistical significance using mixed model analysis with a Bonferroni corrected confidence interval using SPSS (IBM, Chicago USA). For all tested parameters, clot volumes were compared across all flow chambers, with the blood donor ID as the subject variable. The independent variables included: velocity, packing density, distance from the inlet. The dependent variable was the volume of clot as a function of distance. aPTT was used as a covariate. For resistance analysis, the blood donor ID was used as a subject variable, the repeated-measure variable was time, and the independent variable was the condition tested (e.g. velocity or packing density). The dependent variable was set to be ln(resistance) in the SPSS software and aPTT was set as a co-variate. A one-way ANOVA compared the clot volumes at the equivalent distance for the packing density and velocity groups. For path length groups, clot volume for a 1 mm thick segment was compared at 10 mm and 20 mm from the inlet for both devices. Post-hoc tests were completed using Tukey’s test to find the difference between groups. In all cases, a p-value of < 0.05 was regarded as significant.

## 3. Results

### 3.1 Inlet and Baseline Conditions

The donor blood collected in this study was all within normal clinical ranges, albeit slightly on the lower end of normal for hematocrit, hemoglobin, and platelet counts (Table 1Table). The variation between groups was generally small.

**Table 1.** Hematology of each donor group. All results are within normal clinical ranges. N = 6 for each group.

|  | *aPTT (s)* | *PT (s)* | *HCT (%)* | *Hgb (g/dL)* | *PLT (k/uL)* |
| --- | --- | --- | --- | --- | --- |
| Flow Rate | 36.9 ± 3.5 | 8.8 ± 0.3 | 38.8 ± 0.9 | 13.7 ± 0.3 | 237 ± 8 |
| Path Length | 36.0 ± 3.8 | 9.1 ± 0.3 | 39.5 ± 1.9 | 13.4 ± 0.7 | 275 ± 17 |
| Porosity | 41.2 ± 3.8 | 8.4 ± 0.1 | 40.9 ± 1.6 | 13.8 ± 0.4 | 240 ± 21 |

Table 2 contains the baseline, glycerol-water resistances and Reynolds numbers for each test conditionTable. Resistance increases only slightly with packing density and is about 1.68 times greater at a 4 cm path length when compared to a 2 cm path length. Blood flow resistances above these values were thus indicative of clot formation. Reynolds numbers for each flow chamber and condition were calculated using a glycerol-water mix as well, and remained in the clinical range as seen in artificial lungs (Table 2).

**Table 2.** Baseline resistance values for each packing density and path length condition.

|  | **2 CM** |  |  | **4 CM** |
| --- | --- | --- | --- | --- |
| **PACKING DENSITY** | 40% | 50% | 60% | 50% |
| **RESISTANCE**<br>(mmHg/(mL/min)) | 0.333 ± 0.003 | 0.365 ± 0.008 | 0.379 ± 0.005 | 0.615 ± 0.010 |
| **REYNOLDS NUMBER** | 97.4 | 77.9 | 64.9 | 77.9 |
| **ROOT MEAN SQUARE**<br>**SHEAR STRESS (Pa)** | 0.31 | 0.34 | 0.37 | 0.32 |

### 3.2 Effects of Fiber Bundle Design on Clot Formation

#### 3.2.1 General Patterns of Clot Formation

Generally, there was more fibrin deposition in between the fibers and creating bridges, rather than on the surface of individual fibers. On the other hand, platelet deposition was greater directly on the surface of the fibers than the space in between them. These surfaces were likely first covered with proteins, which acted as a nucleation surface for fibrinogen and platelet deposition. Fibrin networks formed in the direction of flow, and subsequently there was less fibrin deposition on the upstream side of the hollow fibers, and more fibrin on the downstream side as seen in Figure 2b. Computational models performed by Zhang et al. have shown that stagnation points exist on both upstream and downstream center lines of the fibers, hence, these are likely sites of clot nucleation.^11^

#### 3.2.2 Effects of Packing Density

Both µCT (Figure 3) and fluorescence microscopy results (Figure 4) indicated greater formation of clot in the densely packed flow chambers than the loosely packed chambers. As seen in the probability maps, clot formed preferentially downstream, near the outlets of the channels (Figure 3a). The more tightly packed bundle had clot filling the proximal regions throughout the length of the chamber. The looser bundle had denser clot but within a smaller distal region, near the outlet. There was also noticeable capture of clot at the inlet of each device, possibly from clot embolizing from upstream locations.

**Figure 3.**
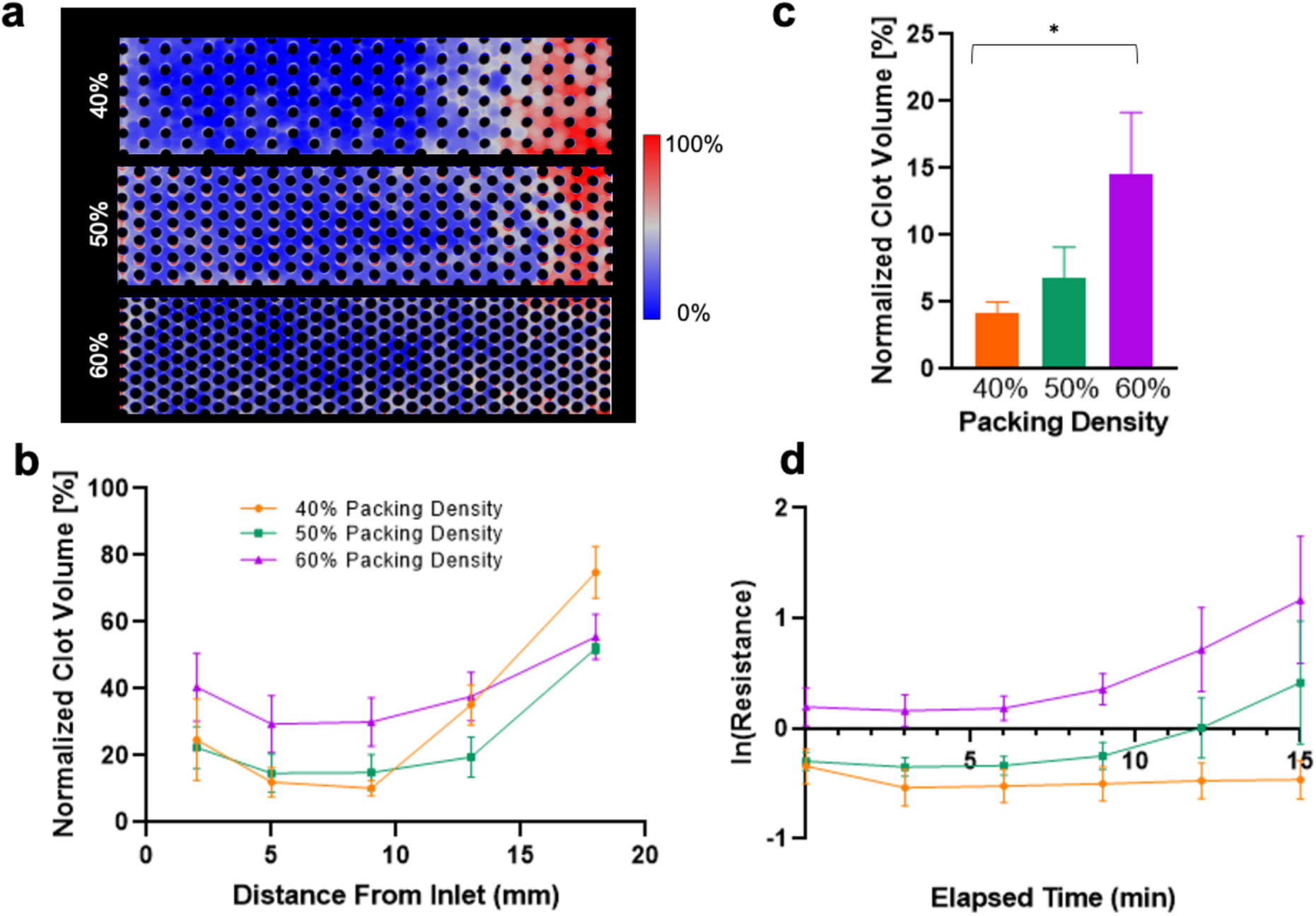
(a) Clot deposition probability maps for 40%, 50%, and 60% packing densities. (b) Clot volume in flow chambers with various packing densities. The 60% packing density device had more clot than the 40% or 50% packing density devices in the entire length of the flow chamber. (c)The clot volume in a 1 mm slice at equivalent distances from the inlet in flow chambers with various packing densities after a 15-minute experiment. (d) The change in resistance in natural log over the 15-minute experiment. A higher packing density correlated with a higher resistance.

**Figure 4.**
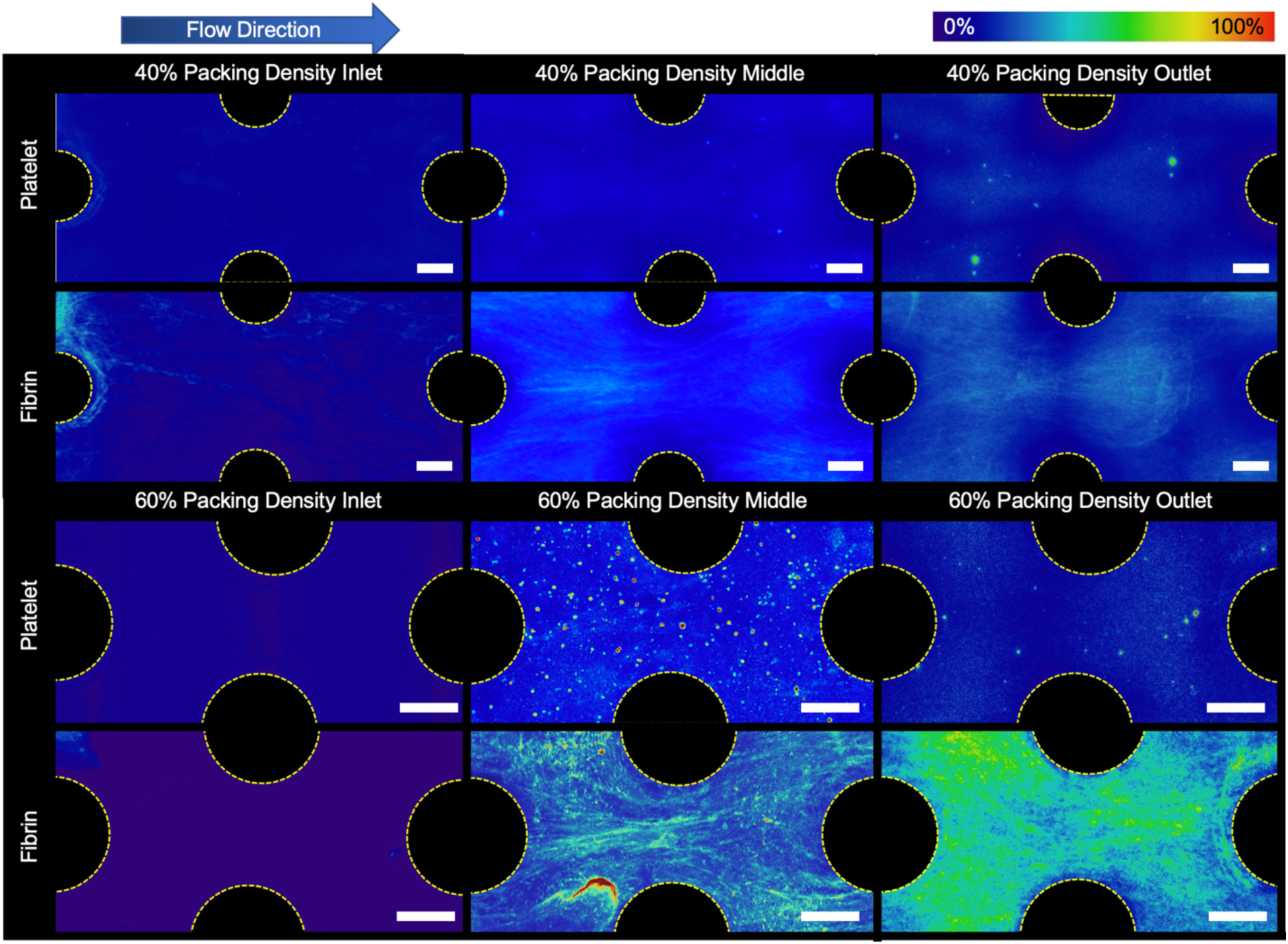
Fluorescently stained platelet and fibrin average deposition in a 40% and 60% packing density flow chamber at the end of the experiment. There was noticeably more fibrin deposited between individual fibers in the 60% packing density devices than in the 40% devices. (Scale bar = 200 µm).

This effect was more prevalent in the 60% packing density flow chambers. The normalized clot volume was significantly higher in the 60% packing density (p = 0.024) in the front half and middle of the flow chamber. (Figure 3b). This is reflected in the quantification of clot at locations in the chamber with equivalent surface areas (Figure 3c). In this case, when accounting for surface activated clot, the 60% packing density flow chamber had more clot than the 40% (p = 0.047). The 40% packing density device also had more clot than the 50% and 60% device at the outlet, were not significant (p = 0.65 and p = 0.24, respectively).

Blood flow resistance measurements confirmed that the rate of clot formation, and thus blood flow occlusion, occurred more rapidly with increasing packing density (Figure 3d). Additionally, the rate of occlusion was found to accelerate at approximately the 8 minute mark in the flow chambers with 50% and 60% packing density chambers, but not in those with 40% packing density. The blood flow resistance in the flow chambers with 60% packing density was found to be significantly greater than in those with 50% (p << 0.001) and 40% packing density (p << 0.001). The resistance of the flow chambers with 50% packing density was also significantly greater than in those with 40% packing density (p < 0.001).

Microscale observations revealed that platelet and fibrin adhesion was lower in the flow chambers with 40% packing density than in those with 60% packing density (Figure 4). Additionally, more fibrin strands formed perpendicular to flow in the flow chambers with 60% packing density, connecting adjacent hollow fibers.

#### 3.2.3 Effects of Blood Flow Velocity

Clot formation increased with lower blood flow velocity in the miniature flow chambers. As can be seen in the probability maps, more clot formed throughout the entire length of the slowest flow chamber as depicted in red and white (Figure 5a). Greater clot density was localized at the outlet of each flow chamber, regardless of velocity. The µCT clot volumes throughout the channels were significantly greater under the lowest flow than the highest flow (p = 0.014) (Figure 5b). At an equvalent distance within each chamber at different flowrates, the clot volume was quantified to account for the effect of surface activation of the coagulation cascade (Figure 4c). At these points, the fastest flow had 2.5 times less clot compared to the medium flow (p = 0.313) and slowest flow (p = 0.213).

**Figure 5.**
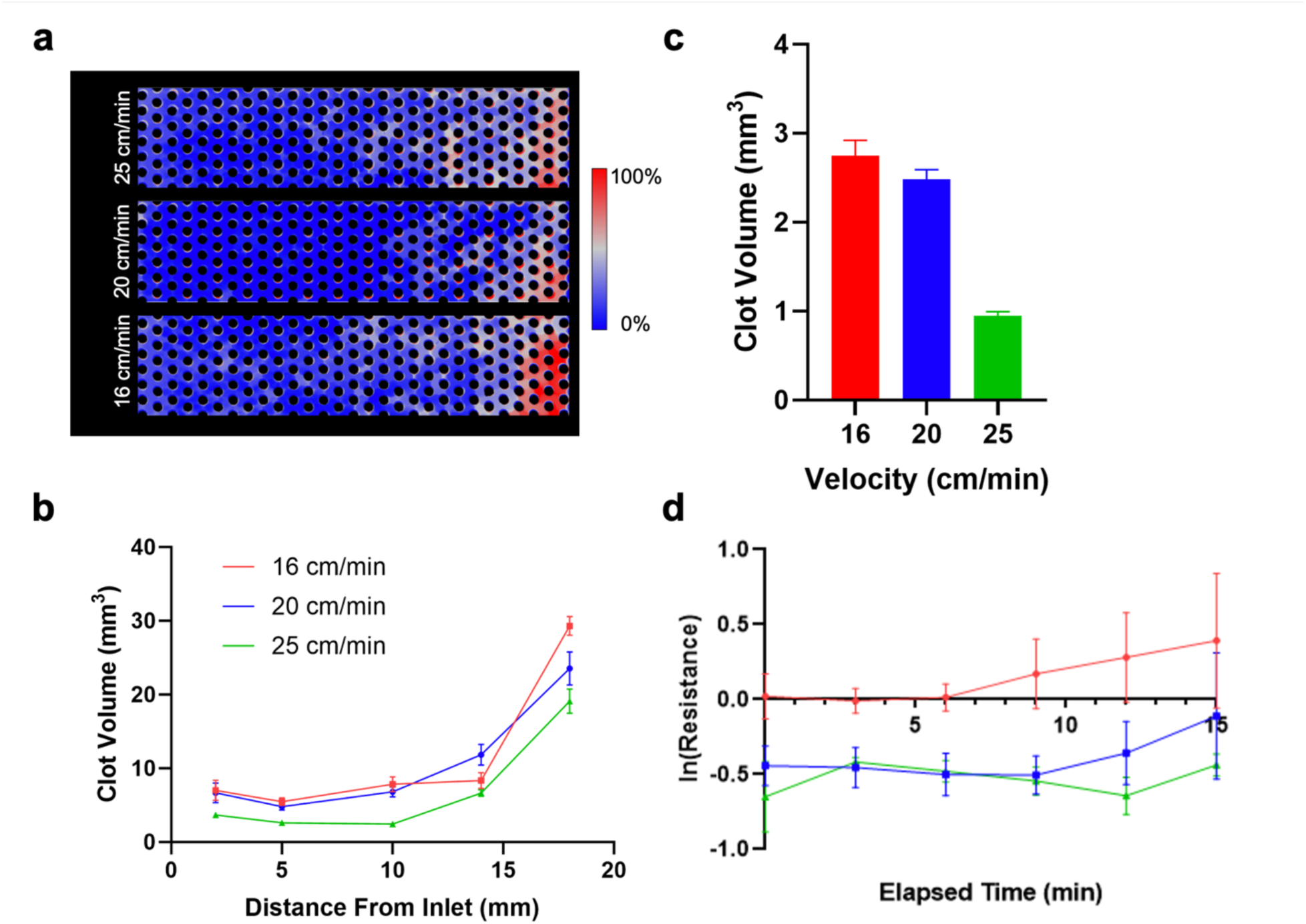
(a) Clot deposition probability maps for 25, 20, and 16 cm/min velocities. (b) Clot volume in flow chambers with various blood flow velocities. (c) The clot volume in a 1 mm slice at equivalent distances from the inlet in flow chambers with various velocities after a 15-minute experiment. (d) The change in resistance in natural log over the 15-minute experiment. A lower flow device was associated with a higher resistance.

Blood flow resistance also followed the µCT results and was significantly greater in the lowest velocity flow chamber compared to medium velocity (p < 0.01) and high velocity (p < 0.001) (Figure 5d). In the medium and high velocity flow chambers, there was a decrease in resistance from 3 to 10-12 minutes and then an increase in resistance thereafter compared to baseline (Table 2). These results indicate that a greater velocity delayed the initiation of clot formation in a hollow fiber bundle.

Confocal microscopy revealed that fibrin networks that form under lower flow rates had thicker strands of fibrin that formed between the hollow fibers in each flow chamber. Additionally, the fibrin network density was also greater with more interconnections between the fibers. As seen in Figure 6, there are qualitatively more fibrin strands surrounding the rods in the low blood flow case at the inlet, middle, and outlet of the flow chamber.

**Figure 6.**
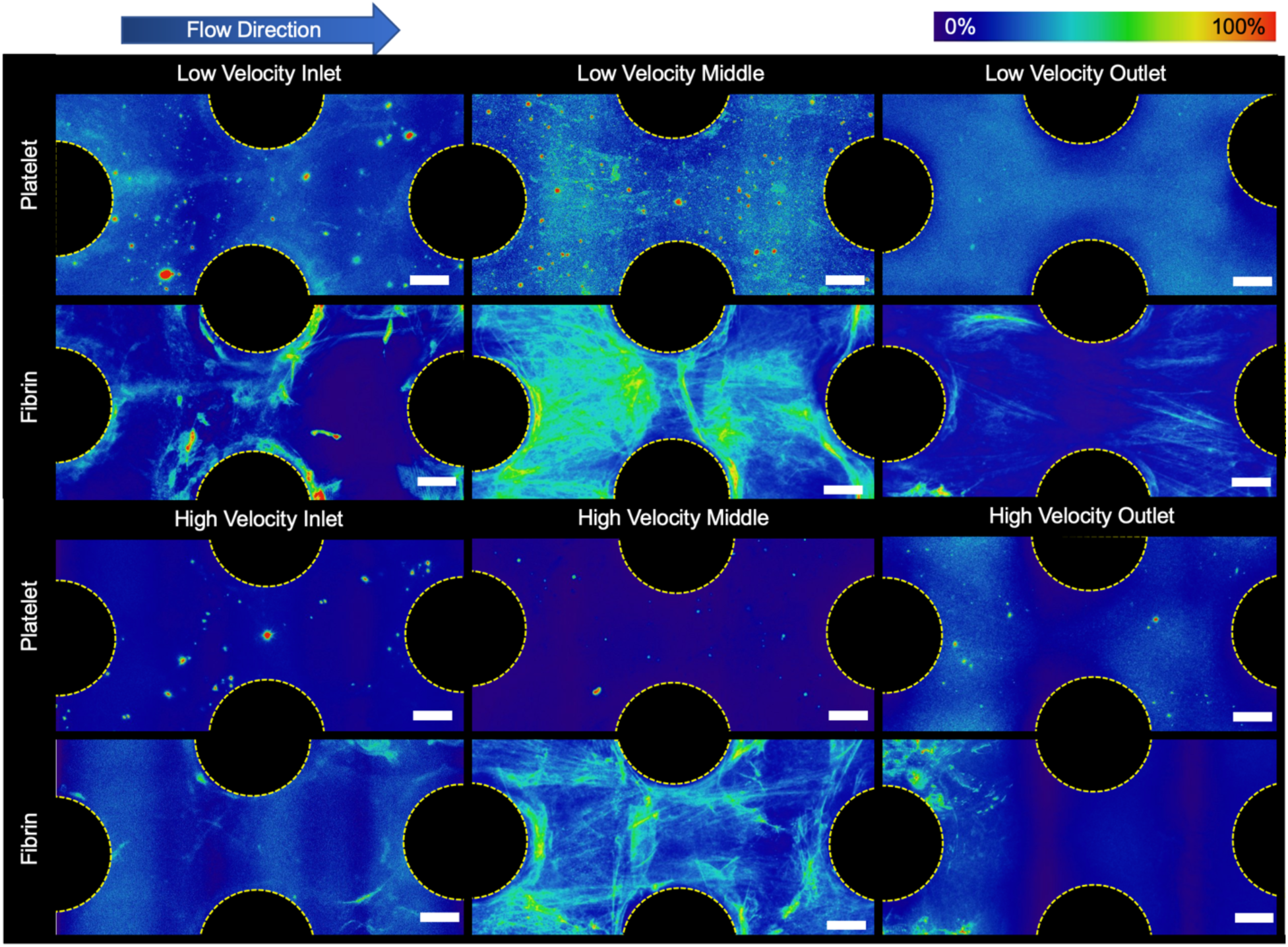
Fluorescently stained platelet and fibrin average deposition in a low (16 cm/min) and high (25 cm/min) velocity blood flow at end of the experiment. There were noticeably more fibrin and platelets deposited between individual fibers in the lower velocity devices. Additionally, there were more distinct trailing strands of fibrin and of captured platelets off the back side of the rods seen in the lower velocity experiments. (Scale bar = 200 µm)

#### 3.2.4 Effects of Path Length

The clot patterns for both 2 cm and 4 cm flow chambers exhibited similar patterns, with the majority of thrombus accumulating at the outlet, while the center of the chamber remained clear. Overall, a shorter path length of 2 cm resulted in a higher volume of deposited clot than in the 4 cm flow chamber. The clot probability map also shows that the shorter flow chamber had more condensed clot through the length of the flow chamber, whereas most of the flow chamber with a 4 cm path length was clear. The clot volume was greatest at the entrance, the first quarter, middle, third quarter, and the downstream end of the 2 cm flow chambers when compared to the 4 cm flow chambers (p = 0.08). (See Figure 7a.)

**Figure 7.**
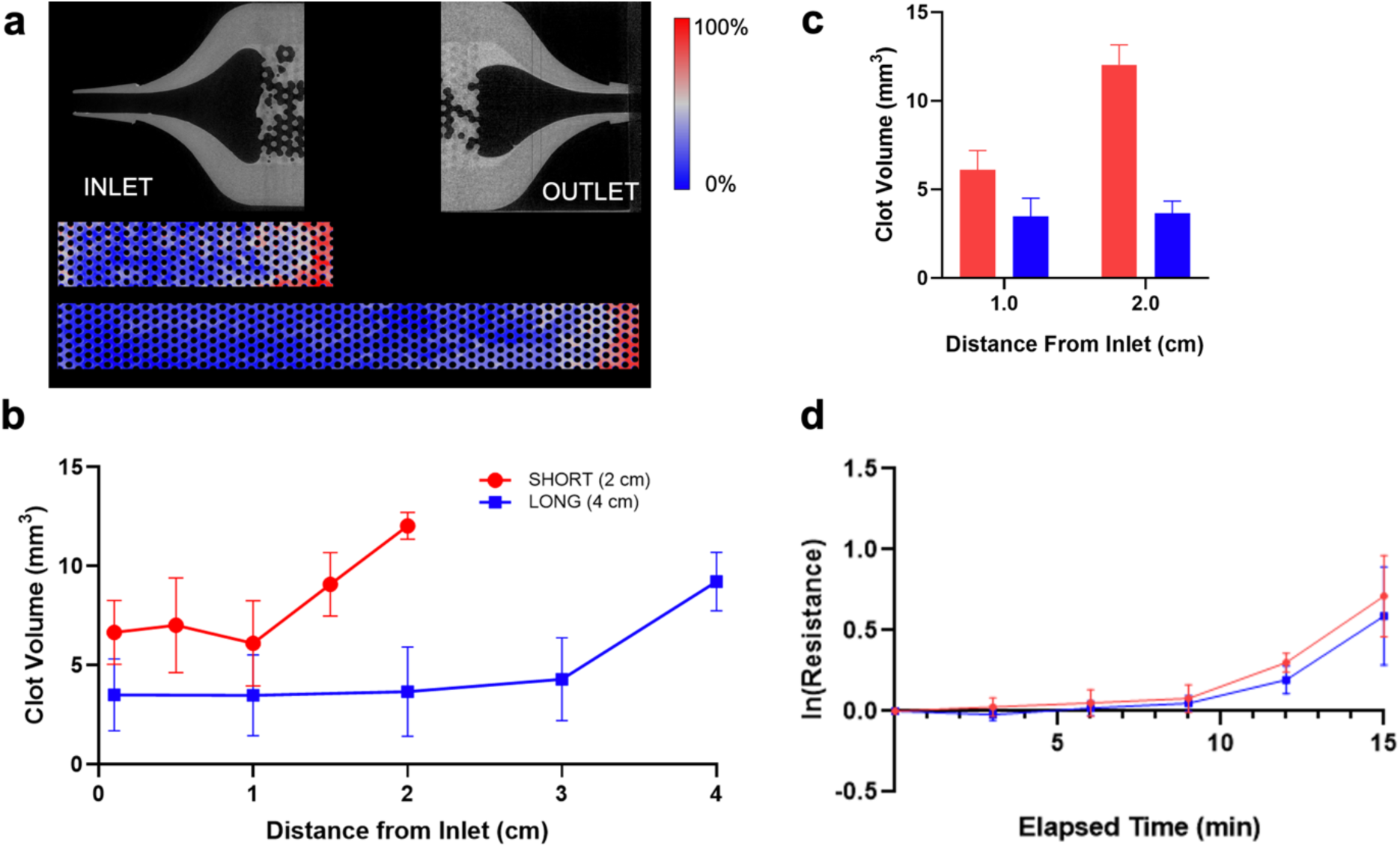
(a) Clot deposition probability maps for 2 and 4 cm path lengths. A representative microCT scan of the inlet and outlet are also shown with the plastic in grey, and the clot in light grey. There is more clot deposited at the outlet. (b) Clot volume in flow chambers with a 4 cm path length had less clot that the shorter 2 cm device. (c)The clot volume in a 1 mm slice at equivalent distances from the inlet in flow chambers with various velocities after a 15-minute experiment. (d) The change in resistance in natural log over the 15-minute experiment. There was no significant difference in resistance between the two path lengths.

Although the clot volume in the 2 cm flow chamber was greater, the blood flow resistance was not significantly different from the 4 cm flow chamber (p = 0.396, Figure 7b). The resistance increased at an increasing rate over 15 minutes for both flow chambers with either 2 cm or 4 cm path lengths in a similar fashion.

To account for the effect of surface area on the two path lengths, the clot volume was also compared at equal distance of 1 cm and 2 cm from the inlet for both flow chambers. (See Figure 7c.) Despite equivalent surface area wetted by blood, there was significantly more clot in the 2 cm flow chambers than in the 4 cm flow chambers at a 2 cm measurement (p = 0.004) but not significantly different at 1 cm and (p = 0.393). This observation suggested that clot formation at the outlet is able to promote formation of clot upstream.

## 4. Discussion

The results from this study indicate that:

1) Clot formation in a microfluidic hollow fiber bundle decreases with decreasing packing density and increasing blood flow velocity,
2) Clot formation at the fiber bundle outlet can significantly increase upstream fiber bundle clot formation, and
3) A longer path length provides more clot-free fiber surface area for gas exchange than a shorter path length due to clot formation at the outlet.

Clot formation decreased with packing density. With a lower packing density, the distance between individual hollow fibers is greater, which impedes the bridging of fibrin strands between adjacent fibers. This results in a less dense fibrin net, which reduces the capture of platelets. This likely also reduces the quantity of red blood cells captured, although not measured here, which ultimately leads to less macro-scale clot formation. Although fiber bundles with a looser design will have prolonged functionality due to lower clot burden, they would provide less oxygenation per total volume due to the lower surface area. Thus low-density fiber bundles would need to be designed with larger path lengths or frontal areas to provide equivalent surface area for gas exchange. Of these two choices, greater frontal area is the least attractive with regards to fiber bundle clot formation, as it reduces blood flow velocity, which increases clot formation. However, the greater path length will also increase resistance, which may induce greater shear stress in the artificial lung and associated blood pump, and in turn may exacerbate platelet activation.^13^

Slower blood velocities through the flow chambers resulted in more clot deposition, as expected due to the lower rates of convection to and from the biomaterial surface.^14^ The intrinsic pathway of the coagulation cascade was activated at the blood-material interface. With slower blood flow, the flow chamber rapidly accumulated more procoagulant factors within the densely-packed, high surface area of the fiber bundle.^15^ Lower velocities may also result in lower shear stresses and will reduce surface washing or enzymatic degradation of a forming clot. In this study, calculated shear stresses were comparable between the studied chambers, and the magnitude was not large enough to shear-induced activation of platelets (Table 2).^16^ Ultimately, low fiber bundle blood flow velocities should be avoided. However, shear stresses above 15 dyne/cm^2^ should also be avoided.

Of note, the majority of clot deposition was consistently found at the outlet of the flow chambers, close to the exit collector, and to a lesser extent at the inlet for all test conditions. This is similar to what is observed clinically.^17^ Artificial lung devices are known to frequently catch thromboemboli at the fiber bundle inlet that originate from procoagulant upstream locations, such as small stagnation regions at the circuit connectors or shear induced platelet aggregates from mechanical pumps. Artificial lungs used for several days to weeks also often have more extensive, occlusive thrombus within their outlet manifold. The outlet commonly has more clot formation due to (1) accumulation of activated clotting factors and activated platelets as blood flows through the fiber bundle and (2) regions of stagnation or recirculation in those regions. In this study, blood almost certainly accumulated procoagulants due to contact system activation, as demonstrated by progressive fibrin formation from inlet to outlet (Figure 4). Additionally, the sudden flow reduction from the wider flow chamber to the 1/16” barb connection at the outlet could have led to regions of slower flowrates. Ultimately, these results suggest that design of the outlet manifold is a neglected factor that may mitigate or exacerbate the extent of clot formation.

Path length of the flow chamber affected the formation of clot to a lesser extent than the other parameters studied. For both the 2 cm and 4 cm flow chambers, a similar level of clot formation formed at the outlet, leading to a comparable increase in blood flow resistance. This suggests that the outlet housing was the dominant cause of clot formation in these devices rather than the path length. In the shorter flow chambers, this clot occluded a greater percentage of the entire flow chamber, which would reduce gas exchange function. Therefore, for full-scale artificial lungs, a longer path length could potentially protect a greater proportion of fibers from occlusive clot forming at the outlet. However, a fiber bundle with a longer path length has more resistance to blood flow. This, in turn would require greater pump speed to generate the required pressure in a an ECMO circuit, leading to increased shear activation of platelets. Thus, the best approach is to carefully design the fiber bundle to avoid stagnation and recirculation and potentially offset the fiber bundle at a distance from the walls of the outlet housing, to avoid clot growing back into the fiber bundle.

Limitations of this study include the use of a miniature flow chamber that was 3D printed with a urethane acrylate – unlike clinical grade artificial lungs that typically employ polymethylpentene gas exchange fibers. While the material selection may change the extent of protein adsorption, and the rate of clot progression, it is not likely to affect the general mechanisms of clot formation caused by all hydrophobic polymer surfaces. This artifact was further mitigated in this study by performing paired comparisons of the independent variables: packing density, length, and velocity. An additional physical limitation was the 3 mm height of these flow chambers, far smaller than the thickness of a full-sized fiber bundle. This increased edge effects but was necessary to quantify clot inside the chamber with different imaging modalities and was mitigated by eliminating areas near the edges from analysis.

Additional artifacts could have been generated due to the artificial surfaces of blood wetted components of the circuit upstream of the flow chamber. However, the same circuit was used in each of the studies to minimize this effect. Nevertheless it would be highly advantageous to confirm the results of this in-vitro study with full-scale artificial lungs having different fiber bundle and outlet housing designs in a large animal model, in which anticoagulation could be titrated to typical of ECMO conditions.

## 5. Conclusions

Multi-scale analysis of clot formation in microchannel devices indicated that fiber bundle clot formation is reduced when there is (1) lower fiber bundle packing density, (2) greater blood flow velocity, and (3) a less procoagulant outlet manifold. These factors reduce fibrin mesh formation, platelet deposition, and bulk clot, particularly at the outlet of the fiber bundle.

## Acknowledgements

R01HL089456

## Supporting Information

**Supplementary Table 1.**
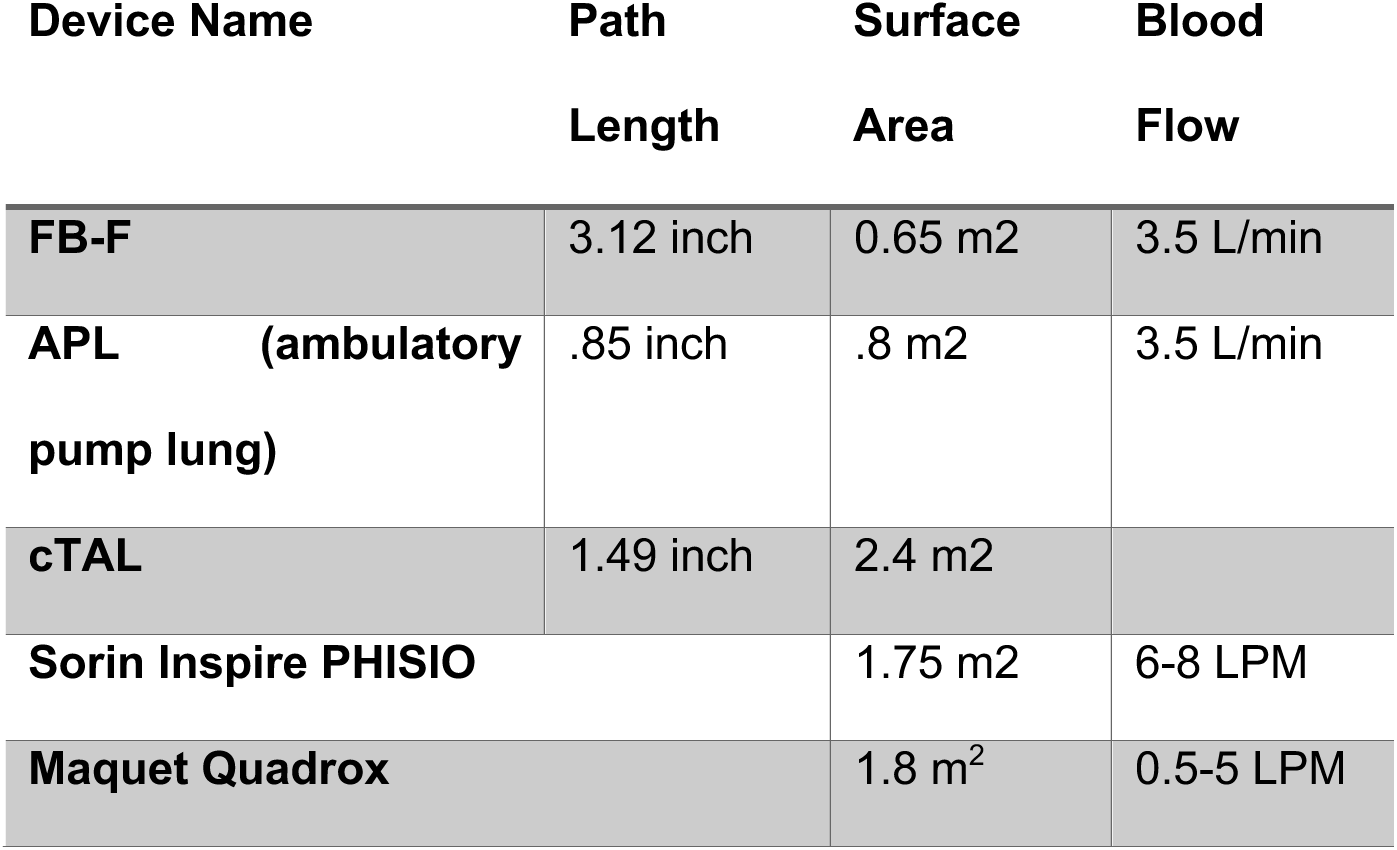

## Supplementary Information 2

`//Open image stack you are interested in that is in binary. Make sure it is saved as a Tiff stack`

`//that has already been cropped and aligned using the 3DSlicer software`

`//`

`//This macro runs through the slices of an already opened TIFF stack to collect the histogram`

`//value for each bin, 0-255, a grayscale 8-bit image`

`run("Clear Results");`

`setOption("ShowRowNumbers", false);`

`for (slice=1; slice<=nSlices; slice++) {`

`“setSlice(slice)”; //sets the slice as the current slice`

`getRawStatistics(n, mean, min, max, std, hist);`

`for (i=0; i<hist.length; i++) {`

`setResult("Value", i, i);`

`setResult("Count"+slice, i, hist[i]);`

`}`

`}`

